# Normalization strategies for lipidome data in cell line panels

**DOI:** 10.1101/2024.07.16.600977

**Authors:** Hanneke Leegwater, Zhengzheng Zhang, Xiaobing Zhang, Thomas Hankemeier, Amy C. Harms, Annelien J. M. Zweemer, Sylvia E. Le Dévédec, Alida Kindt

## Abstract

Sample collection can significantly affect measurements of relative lipid concentrations in cell line panels, hiding intrinsic biological properties of interest between cell lines. Most quality control steps in lipidomic data analysis focus on controlling technical variation. Correcting for the total amount of biological material remains an additional challenge for cell line panels. Here, we investigated how we can normalize lipidomic data acquired from multiple cell lines to correct for differences in sample biomass.

We studied how commonly used data normalization and transformation steps during analysis influenced the resulting lipid data distributions. We compared normalization by biological properties such as cell count or total protein concentration, to statistical and data-based approaches, such as median, mean, or probabilistic quotient-based normalization and used intraclass correlation to estimate how similarity between replicates changed after normalization.

Normalizing lipidomic data by cell count improved similarity between replicates, but only for a study with cell lines with similar morphological phenotypes. For cell line panels with multiple morphologies collected over a longer time, neither cell count nor protein concentration was sufficient to increase the similarity of lipid abundances between replicates of the same cell line. Data-based normalizations increased these similarities, but also created artifacts in the data caused by a bias towards the large and variable lipid class of triglycerides. This artifact was reduced by normalizing for the abundance of only structural lipids. We conclude that there is a delicate balance between improving the similarity between replicates and avoiding artifacts in lipidomic data and emphasize the importance of an appropriate normalization strategy in studying biological phenomena using lipidomics.

## Introduction

The lipidome, *i.e.,* the total set of lipids in a cell, is a subgroup of the metabolome comprising fats, oils, and other related substances. Lipids are classified based on their structural characteristics, with distinct lipid classes performing distinct biological roles. These functions encompass vital aspects including energy storage, membrane structure and fluidity, signaling processes, and overall cellular function (Han, 2016). Metabolic processes related to lipid metabolism are altered in cancer cells, promoting their growth, proliferation, and migration (Beloribi-Djefaflia et al., 2016). All these are properties of a more aggressive phenotype (Broadfield et al., 2021). Thus, analyzing the lipidome of cancer cells can provide insight into the malignancy of specific cancer phenotypes and information on the metabolic state of the system being investigated.

It is essential to get an accurate measure of the abundance of individual lipid species to understand the underlying biology correctly. Advances in mass spectrometry (MS) have accelerated research performed in this field (Blanksby & Mitchell, 2010; Han et al., 2012). However, technical variation before or during the measurements can influence final metabolite concentrations. This variation may occur at any point in the experimental setup, starting with the collection of cell pellets. Sample preparation influences which lipids are extracted (Sethi & Brietzke, 2017) and how the lipids are measured and quantified (Köfeler et al., 2021). Many quality control steps are employed during lipidomic experiments that focus on removing technical variation, including the use of internal standards (Fan et al., 2019; Vaz et al., 2015), removal of background signal (Köfeler et al., 2021) and batch correction (Züllig et al., 2020).

However, these quality control steps do not correct for variation in sampling. Overall differences in cell size, different cell numbers due to different proliferation rates or other experimental factors during the collection of cell pellets can create unwanted variation. It is essential to normalize data to be able to compare metabolite abundances between cell lines in a panel. This can be performed using biology- or data-based approaches. Examples of biology-based normalizations are standardization to cell count, concentration of a molecule such as DNA or protein, stable ion representatives (Chao et al., 2017) or quality control metabolites, which are metabolites that are expected not to change during an experiment (Livera et al., 2015). Each of these methods has potential issues. These approaches assume that the cells have similar sizes, morphological phenotypes, e.g., only mesenchymal or only epithelial morphologies(Chao et al., 2017), or assume the same ploidy (Muschet et al., 2016; Quinton et al., 2021). Imaging data based on cell confluency could be used (Ortmayr et al., 2019), but this technology is not always available. Thus, correcting for the total amount of biological material remains a challenge in cell line panels (Livera et al., 2015).

As an alternative approach to biology-based normalizations, data-based normalizations assume an overall shared concentration over all metabolites in a sample. Normalization by median signal assumes that cell lines have the same median lipid abundance, while normalizing using mean or total sum approaches assume that the total amount of lipid abundance is the same in each sample. This is more sensitive to extreme positive values than median normalization. Another often used technique in metabolomics is the probabilistic quotient normalization (PQN), which calculates an overall dilution factor for each sample when compared to a reference sample within the dataset, with the assumption that the net changes in the data are zero (Dieterle et al., 2006). For metabolomics data, this would mean that all samples have similar overall distributions.

Many tools exist for data normalization, for example metaboAnalyst (Pang et al., 2022) or lipidr (Mohamed et al., 2020), and for assessing normalization quality, such as Normalyzer (Chawade et al., 2014) and NOREVA (Li et al., 2017). However, Normalyzer misses the option to allow a biological measure for normalization, while NOREVA does not allow easy comparisons between two normalization strategies of interest. In addition, it is not always clear which normalization strategy is best for a specific situation when using metaboAnalyst or lipidr.

In this study, we investigate normalization techniques for a lipidomics dataset acquired on a panel of cancer cells with a wide variety of sizes, proliferation rates and morphologies, resulting in suggestions and possible caveats for fellow lipidomics scientists.

## Methods

### Cell culture

A total of 52 breast cancer cell lines with various morphologies were obtained from the Erasmus MC, the Netherlands, and stored frozen in liquid nitrogen before use. Cells were passaged for a limited time to prevent population and genetic drift. All cell lines were cultured in RPMI 1640 with 10% fetal bovine serum, 25 IU/mL penicillin and 25 µg/mL streptomycin at 37°C, 5% CO_2_. Cells were seeded in 10 cm dishes in 10 mL media. Media was refreshed 24 hours after seeding cells, with a starting density that aimed for 80% confluency at the end of the experiment and cells were grown for 3 days. For each replicate of a cell line, a new vial of the same passage was thawed and split twice using trypsin before use in experiments. Replicate experiments were performed on different days.

### Cell line panel morphology assessment

Cell morphological phenotypes were obtained from Rogkoti *et al*. (Rogkoti et al., n.d.). For missing cell lines, we defined the morphology as epithelial-like, mesenchymal-like or round for adherent cell lines or suspension for non-adherent cell lines, based on available images from the ATCC (ATCC, n.d.) and descriptions of Cell Model Passports (van der Meer et al., 2019). Cell lines without identifiable morphologies were left as unknown (**Supplementary Table 1**).

### Drug treatment

To study a dataset with less expected biological variation, we selected three cell lines with a mesenchymal morphology, MDA-MB-231, Hs578T and HCC38, and seeded 1.5 million, 0.4 million or 0.4 million cells in 10 cm dishes 24 hours before treatment. On T=0, media was refreshed, and 10 mM 2-deoxy-D-glucose (2DG) in dimethyl sulfoxide (DMSO) or DMSO control was added to each treatment or control dish. After 0, 6, 24 or 48 hours, one treated and one control dish per cell line were collected as described below. This experiment was performed twice.

### Sample collection, protein concentration and cell count measurements

No pellets were collected for one replicate of two cell lines (BT483 and HCC1569), due to slow cell growth. For each dish, media was removed and cells were washed with PBS, detached using trypsin, and resuspended in 5 mL RPMI 1640. A 500 µL sample was taken for protein concentration determination and a 10 µL sample was used to count cells using a TC20 automated cell counter (Bio-Rad). The remainder was centrifuged for 1000 rpm, 3 minutes at 4 °C to pellet the cells. Pellets were washed twice using ice-cold saline solution (0.9% w:v) and cell pellets were stored at -80 °C until further sample processing for lipidomics analysis.

Protein concentration was determined using a slightly modified bicinchoninic acid (BCA) assay. 500 µL cell suspension was centrifuged and the pellet was resuspended in 200 µL RIPA lysis buffer (10g/L DOC, 50 mM Tris pH 7.5, 150 mM NaCl, 0.1% SDS, 1% NP40, 2 mM EDTA) with a protease inhibitor cocktail (Sigma-Aldrich, p8340-ML) and stored at -20 °C until further use. 1 µL lysed sample was mixed with 250 µL BCA solution (solution A: solution B 50:1, Thermo Fisher Scientific, 23225), incubated for 45 min at 37 °C and measured at 562 nm using a plate reader. Concentrations were calculated using a calibration line ranging from 0-6 mg bovine serum albumin (Thermo Fisher Scientific) in RIPA lysis buffer.

## Lipidomics data collection

### Sample preparation for mass spectrometry

Lipid extraction was performed using a methyl tert-butyl ether (MTBE) method described previously(Zhang et al., 2023). Briefly, 34 µL of internal standards (IS) mix with one lipid per class was added to cell pellets and vortexed. 231 μL of methanol (MeOH) and 770 μL of MTBE were added to this mixture. The sample was incubated at room temperature on an orbital shaker for one hour, followed by the addition of 192.5 μL of water, making the final ratio MTBE:MeOH:Water (10:3:2.5, v/v/v). The mixture was again incubated at room temperature for 10 minutes and then centrifuged at 15800 rcf for 10 minutes. A volume of 520 µL of upper layer was collected and dried in a vacuum concentrator followed by reconstitution in 200 µL of acetonitrile:methanol (3:7). This mixture was vortexed and centrifuged for 10 minutes. The supernatant was collected for LC-MS analysis.

The batch design included solvent blanks, procedure blanks (with IS), cell study samples and quality control (QC) samples. These QC samples were a pool of all the study cell line samples and were analyzed at regular intervals in the study batch to determine the method’s performance. Four samples, (one replicate each for the cell lines EVSA-T, MDA-MB-468, MDA-MB-361 and MDA-MB-175) were lost during LC-MS sample preprocessing. In total, lipid samples for 46 breast cancer cell lines with 3 replicates and for 6 cell lines with 2 replicates were distributed randomly over two batches for LC-MS measurements, and treatment samples were measured in a third batch.

### Lipid detection and relative quantification

A QTRAP 6500+ (AB Sciex) coupled to an Exion LC AD (AB Sciex) was used for targeted lipid profiling to obtain lipid information, including retention time (t_R_) and MS/MS fragmentation spectra as described previously(Zhang et al., 2023). The column used for the separation was a Luna amino column (100 mm × 2 mm, 3 μm, Phenomenex). The mobile phase A and B were 1mM ammonium acetate in chloroform: acetonitrile (1:9) and 1mM ammonium acetate in acetonitrile: water (1:1). Three injections were made to accommodate all the multiple reaction monitoring (MRM) transitions of the targeted lipid features. The first injection measured sphingomyelins (SM), ceramides (Cer) and ceramide subclasses (dhCer, HexCer, LacCer, and an unknown sphingolipid), and diglycerides (DG) in positive mode. The second injection measured phospholipids (PS, PI, PG, PC), lysophospholipids (LPS, LPI, LPG, LPC) and bis(monoacylglycerol)phosphates (BMP) in negative mode. The structural isomers PG and BMP eluted at the same retention time and could not be distinguished; hence, they were reported as PG/BMP. The third injection measured triglycerides (TG) and phosphatidylethanolamines (PE) using switching polarity mode. The injection volumes were 5 μL for the first and second acquisition run and 1 μL for the third run. The column temperature was kept at 35 °C. The injector needle was washed with isopropanol:water:dichloromethane (94:5:1, v/v/v) after each injection.

MS/MS experiments were done on a QTRAP 6500+ mass spectrometer with a Turbo V electrospray ionization source (AB Sciex). The declustering potential (DP) and collision energy of lipid transitions were optimized to obtain the highest response for a mixture of lipid standards. The information about the precursor ion (Q1), characteristic product ion (Q3), lipid name (ID), optimized DP, and collision energy of all the lipid features were incorporated into the MS acquisition method before screening in samples. The parameters of the QTRAP 6500+ mass spectrometer were as follows: curtain gas (N2) was 20psi; collision gas (N2) kept at medium, electrospray voltage was 5500V and -4500V for positive and negative mode, respectively; source temperature was 400 °C, GS1 and GS2 were 30 and 35 psi, respectively. Scheduled MRM (sMRM) was used for data acquisition for targeted analysis. The total scan time was 0.5 sec. The lipids detected by the UHPLC-QTRAP based lipidomics profiling analysis were processed using AB Sciex OS (version 2.1.6, AB SCIEX).

### Internal standard correction and batch correction

Peak areas for lipid abundances were corrected for variation caused by lipid extraction, ion suppression and run day using internal standards and QC samples using an in-house quality control tool. Briefly, peak areas for each detected lipid were divided by the peak area of the internal standard for that lipid class to correct for lipid-specific and sample preparation-specific variation, creating relative ratios. Between batch correction was performed using median normalization (van der Kloet et al., 2009). Compounds with more than 30% relative standard deviation of the QC and a background signal of more than 40% were removed from further analysis. After initial data processing and quality control, relative peak abundances were obtained for 134 lipids in positive mode, 281 in negative mode and 455 in switching polarity mode.

### Data normalization, transformation, and scaling

Datasets were preprocessed in R v4.3.0 (R Core Team, 2023) for normalization and comparisons. The dplyr package v1.1.4 (Wickham et al., 2023) within the tidyverse framework v2.0.0 (Wickham et al., 2019) was used to merge datasets as needed. Dividing samples by the median or total sum of the signal per sample, or by protein concentration or cell number was performed using base R. PQN normalization was performed as described by Dieterle *et al*. (Dieterle et al., 2006) with the median of all samples as a reference. Log_2_ and cube root transformation was performed using base R. Autoscaling was performed with the scale function of base R.

### Methods to identify the best normalization strategy

To find a method that reduces the effect between replicates, we used a mix of statistical and data visualization tools described below.

### Intraclass correlations and Pearson correlations

The irr R package v0.84.1, based on the ICC package (Gamer et al., 2019), was used to calculate an intraclass correlation (ICC) between samples and replicates for each lipid in a dataset, with a two-way model, agreement, and a single unit. Intuitively, the ICC can be interpreted as a number between 0 and 1, in which 1 is best, meaning that replicates are identical for a single lipid (see Formula 1). A lower value means more variation between replicates for a lipid; ICC is 0.5 if the variation between replicates and the variation between all samples are similar, and below 0.5 if there is little agreement between replicates (Koo & Li, 2016). For a thorough explanation of the mathematics, we refer to Koo & Li (Koo & Li, 2016).

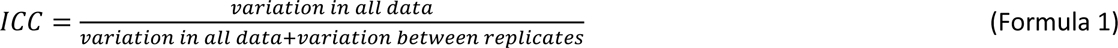

All other correlations were Spearman or Pearson correlations, calculated using the cor function in the stats package in base R. These are reported using the lowercase r for Pearson or r_s_ for Spearman.

### Hierarchical clustering

Hierarchical clustering was performed with Euclidean distances and complete linkage using the pheatmap package v1.0.12 (Kolde, 2019).

### Data visualization

To assess the quality of normalization, density distribution of lipids per group of sample replicates, and boxplots of intraclass correlations per lipid per group per sample were plotted using the ggplot2 package v3.4.4 (Wickham, 2016) and cowplot v1.1.1 (Wilke, 2020). Density plots were based on code from the normalyzer R package (Chawade et al., 2014). Venn diagrams were drawn using ggvenn v0.1.10 (Yan, 2023).

### Assessment of normalization quality

Principal component analysis (PCA) was performed using the PCA function in the mixOmics R package v6.24.0 (Rohart et al., 2017) using autoscaled lipid abundances. Differential lipid analysis was performed after averaging replicates for cell lines, using limma v3.56.2 (Ritchie et al., 2015) on either log_2_ or cube root transformed datasets without scaling after PQN normalization.

### Averaging per lipid class

The average abundance for each lipid class was calculated using autoscaled, centered lipid abundances, which have an average of 0 over all samples. Values were summed for all lipids per class.

## Data and code availability statement

Metabolomics data have been deposited to the EMBL-EBI MetaboLights (Yurekten et al., 2024) with the identifier MTBLS9493 and are accessible at https://www.ebi.ac.uk/metabolights/MTBLS9493. All code to reproduce this analysis can be found at https://zenodo.org/doi/10.5281/zenodo.10635384.

## Results

### Main variation in data before normalization is linked to total signal

This study aims to identify differences in the lipid composition of breast cancer cells with different morphologies. To achieve this, the relative abundances of 870 lipids covering 22 lipid classes are quantified in a breast cancer panel of 50 cell lines with various morphologies grown over 3 days, aiming at 80% confluency at the point of harvesting.

To identify the main variations in lipid abundances, a PCA after log_2_ transformation and autoscaling is performed. This shows a small morphology-related effect (**Figure 1A**), but replicates of the same cell line do not cluster close together regardless of their morphology (**Figure 1B**). However, the total median amount of signal per sample correlates strongly with the first principal component (**Figure 1C, D,** r_s_ = -0.99).

**Figure 1.**
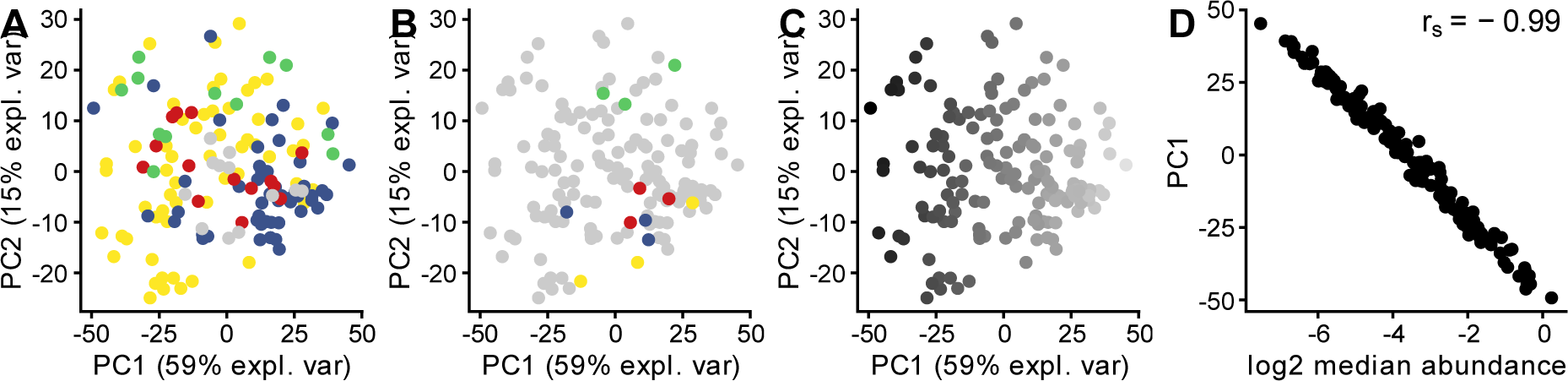
Variation in data is mostly influenced by median signal in a sample. A PCA was calculated using log2 transformed, centered, and scaled data. (A) Morphology was not the basis for the observed separation in the PCA; epithelial-like (yellow), mesenchymal-like (blue), round (red), suspension (green), and unknown morphology (grey). (B) Example cell lines highlighted to show that replicates do not cluster. (C) Median detected signal for the sample drives separation in PC1, which explained 59% of total variation (legend: black to white = high to low log2 median abundance per sample). (D) PC1 and log2 median signal per sample showed a strong negative correlation (rs=-0.99).

### Data-based normalizations outperform biology-based normalization for cell lines with different morphologies

To find a normalization approach that reduces variation between replicates as much as possible, we compared five strategies that can be categorized by their approaches, i.e., biology- or data-based normalizations. The former utilizes the measurement of a biological entity, such as the concentration of protein or DNA, or cell count, with the underlying assumption that the variable represents a proportional amount of assessed material in the sample. The latter utilizes properties of the data for normalization, *e.g.*, the sum or median for a sample, or the overall shape of the distribution, such as during probabilistic quotient normalization (PQN). Intraclass correlations (ICC) per individual lipid are calculated for all samples, where a higher ICC indicates greater similarity between all replicates for a cell line.

When investigating all cell lines, it can be shown that neither cell count nor protein concentration normalization improves the similarity between the replicates. Protein normalization even worsens the ICC between replicates (**Figure 2A**). In contrast, all data-based methods improved the ICC between replicates (**Figure 2A**). Density distributions of all samples show less overlap of distributions either before normalization (**Figure 2B**) or after biology-based normalization (**Figure 2C**) than after data-based normalization (**Figure 2D**).

**Figure 2.**
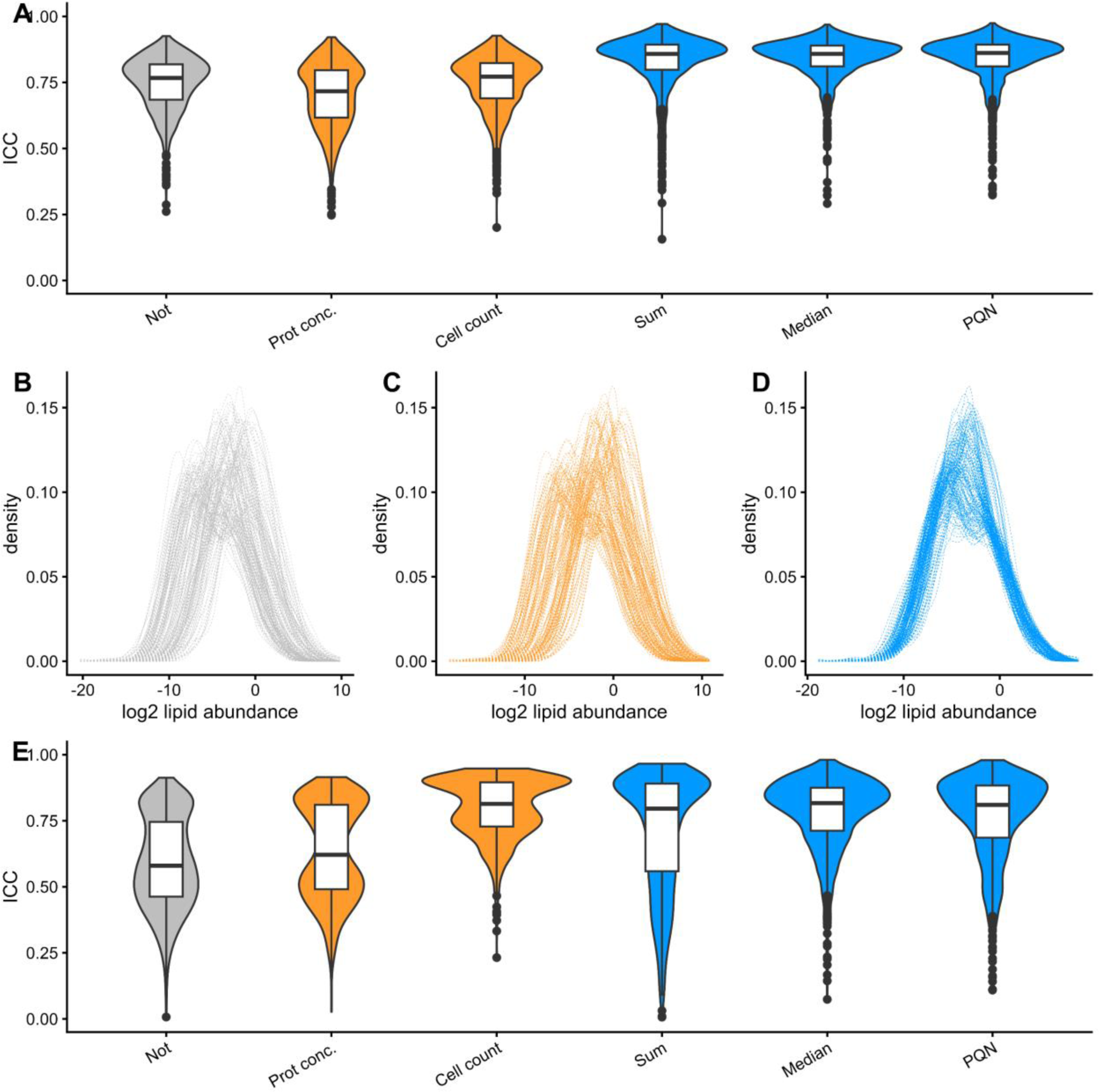
Data-based normalizations improve similarity between replicates. (A) Distributions of intraclass correlations per lipid are calculated before and after normalization for the panel of cell lines. (B-D) Lipid density distributions per sample (B) before normalization, (C) after normalization by protein concentration or (D) after PQN normalization for the cell line panel. (E) Distributions of intraclass correlations per lipid are calculated before and after normalization for cell lines with similar morphologies.

To understand if the observed problems with biology-based normalization are due to the diverse morphologies in the dataset, three cell lines with a similar mesenchymal morphology were investigated further. For each cell line, two replicates were treated with 2DG or a DMSO control and collected at four time points, resulting in 42 samples. In agreement with previous results, normalization by protein concentration does not increase ICC. However, cell count and normalization by sum, median or PQN in the cell line panel show similar improvements (**Figure 2E**). Particularly, PQN and median normalization result in lipids with high ICC values in this dataset (PQN: 636, median: 668, sum: 536, cell count: 711 lipids with ICC > 0.7). Based on this analysis, we conclude that biology-based measures are insufficient to normalize our lipidomic data when cell lines have different morphologies.

### Cell count and protein concentration are inappropriate for lipid abundance normalization

If cell count and protein concentration were effective normalization methods, similar lipid profiles could be expected for cell line replicates with similar protein concentrations or cell counts. However, replicates of cell lines with comparable protein concentrations (T47D=0.40, 0.39, and 0.38 µg/µL, SUM149PT=0.49, 0.47, and 0.48 µg/µL) show different overall lipid distributions (**Figure 3A**). In contrast, cell lines with the largest difference in protein concentration per replicate (OCUB-F=0.39, 0.33 and 0.88 µg/µL, MCF7=0.70, 0.61, 0.95 µg/µL.) have more comparable lipid distributions (**Figure 3B**).

**Figure 3.**
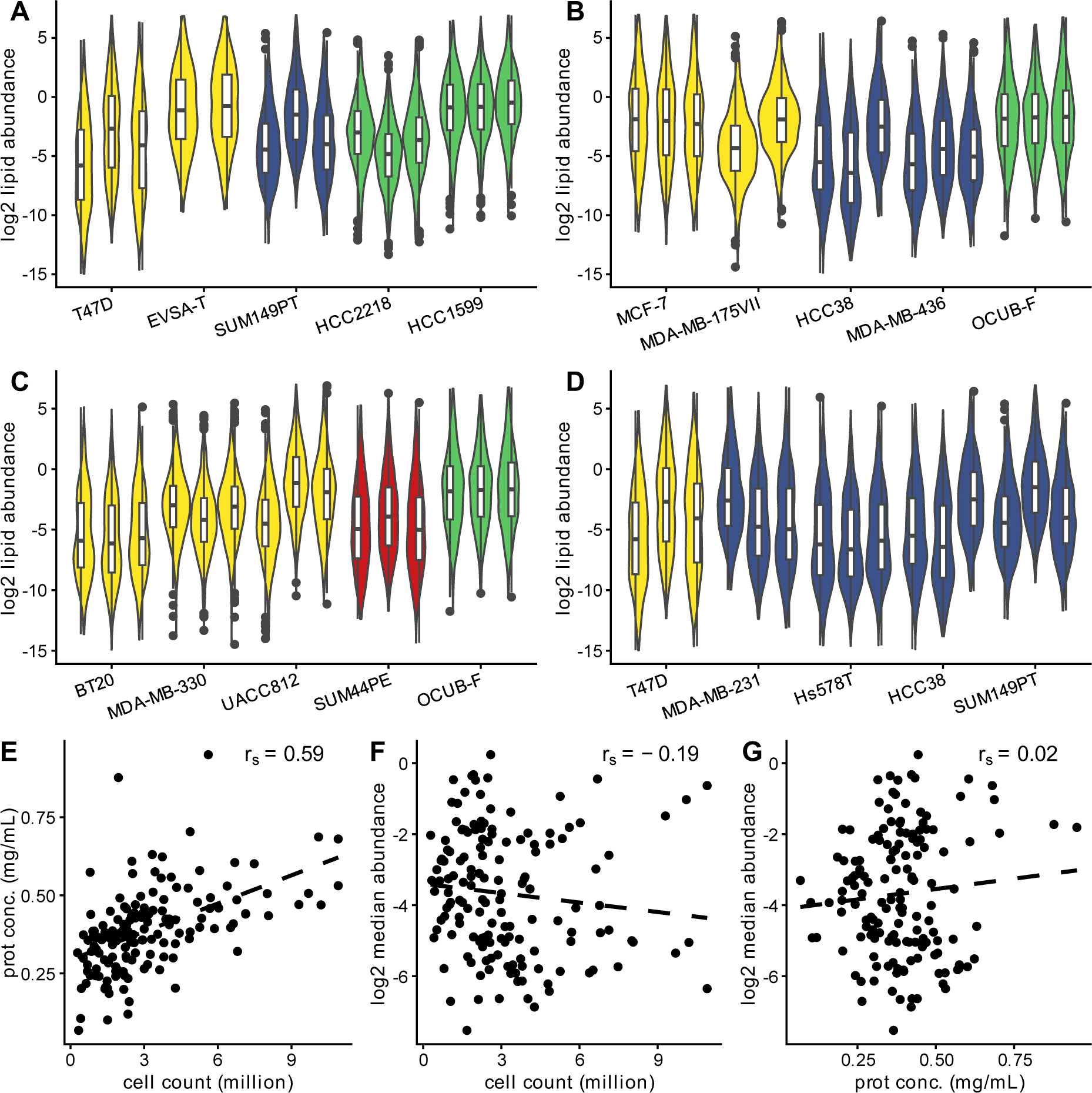
Lipid abundance distributions are not directly related to total protein concentration or cell count. (A-B) Distribution of lipid abundances for 5 cell lines where replicates have (A) the most similar total protein concentration with a standard deviation (sd) 0.00 -0.01 µg/µL, or (B) most different total protein concentrations with a sd 0.15 - 0.30 µg/µL. (C-D) Distributions for cell counts with (C) minimum sd of 0.11 to 0.18 million cells or (D) max sd of 1.7 to 2.1 million cells. Distributions are colored per cell morphology with epithelial-like (yellow), mesenchymal-like (blue), round (red), or suspension (green) morphology. (E-G) Correlation between (E) the two biology-based methods or (F) between cell count or (G) protein concentration and log_2_ median sample abundance. r_s_: Spearman correlation coefficient.

A similar mismatch between total lipid signal and replicate cell count is observed. Replicates of UACC812 with similar cell numbers show a different lipid distribution (UACC812 = 1.3, 1.6 and 1.2 million, **Figure 3C**) and distributions for Hs578T are similar even though they have different cell count values (Hs578T = 2.9, 6.4 and 4.8 million, **Figure 3D**).

Protein concentration and cell count correlate (r_s_ = 0.59, **Figure 3E**), but neither correlates with the median signal per sample (**Figure 3F-G**, r_s_ = -0.19 and r_s_ = 0.02). Thus, neither protein concentration nor cell count are appropriate to correct the variation in lipid abundance in this dataset.

### Lipid classes scale differently with total lipid amount per sample

The assumption of the data-based normalizations evaluated in this study is that replicates of the same cell line have similar overall lipid distributions. If that assumption is valid for this cell line panel, lipid classes should show shared abundance patterns due to the dilution factor of a sample, next to individual cell line or morphology-dependent differences. Phospholipid class averages correlate strongly before normalization (r = 0.85-0.95) and all correlate well with sphingomyelin (r = 0.75-0.8), but a lower correlation is observed between triglycerides and structural lipid classes (r = 0.46-0.65). In general, structural lipids correlate with each other more than lipids that are components of lipid droplets or are involved in signaling (**Figure 4A**).

**Figure 4.**
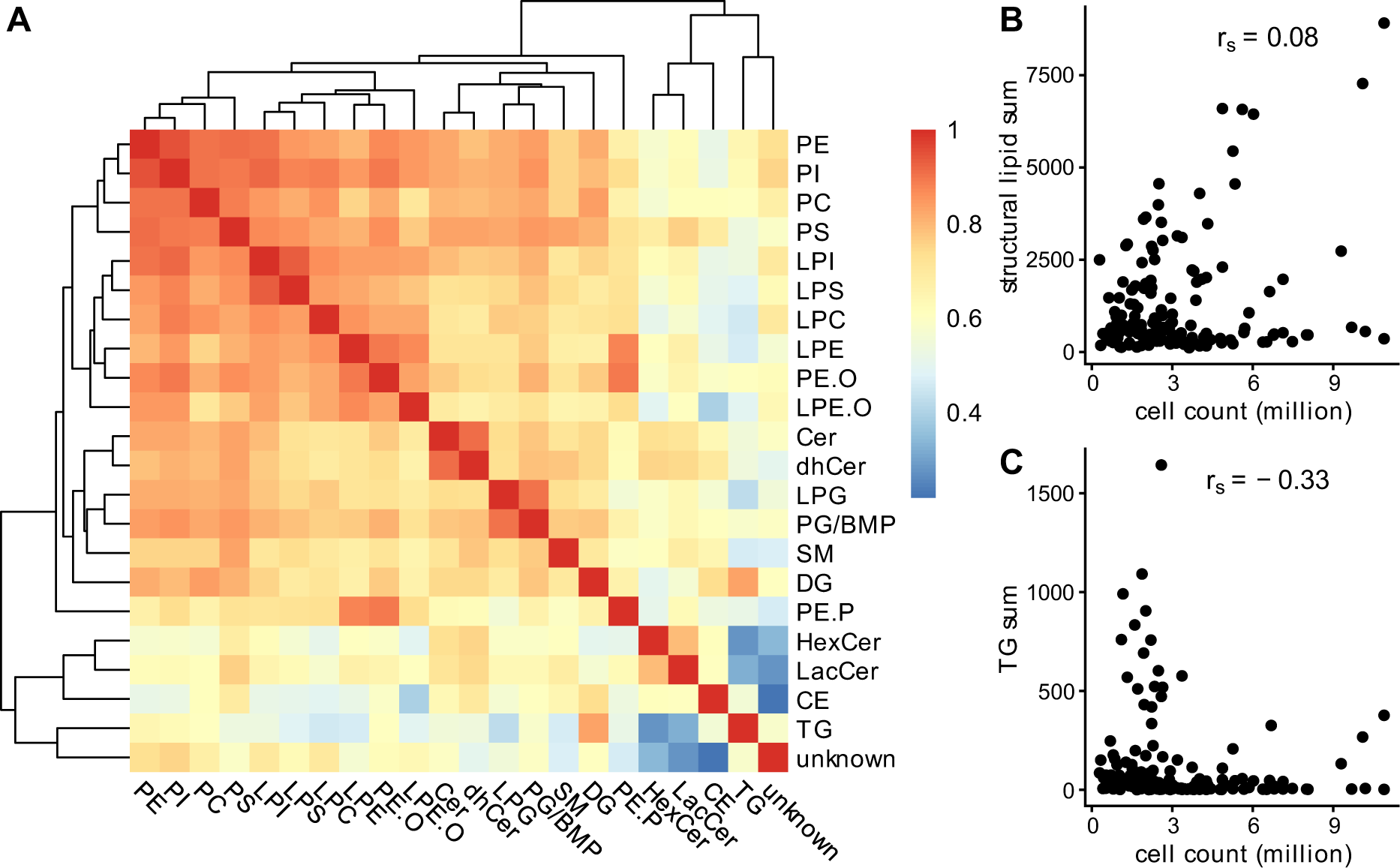
Correlation between lipid classes before normalization shows an overall agreement for structural lipids, but not for triglycerides and some signaling lipids. (A) Correlation between lipid classes. Lipid abundances are log_2_ transformed, scaled, and centered. Class averages are calculated for cell line replicates before normalization, and Pearson correlation is used to correlate class averages over all samples. Rows and columns are clustered using hierarchical clustering with Euclidean distances and complete linkage. (B-C) sum of (B) all structural lipids or (C) triglycerides as a function of cell count. Lipids abundances are not transformed or scaled. r_s_: Spearman correlation coefficient.

We investigated whether this difference between structural lipids and triglycerides explains the low correlation between cell count and lipidomic signal. Although there is no overall correlation between cell count or protein concentration and the sum of structural lipids before transformation (r_s_ = 0.08 and r_s_ = 0.22), there appear to be two populations of samples: samples for which the sum of all structural lipid species scales with cell count and for which it does not (**Figure 4B, Supplementary Figure 1A for protein concentration**). These two populations are not observed for triglycerides (r_s_ = - 0.33, **Figure 4C, r_s_ = -0.09, Supplementary Figure 1B for protein concentration**).

This suggests that the assumption needed for data-based normalization, similar lipid profiles over all cell lines, is not valid due to biological and functional differences of the lipids.

### PQN normalization can create artifacts

Different distributions, abundances, and correlations between classes over the samples can influence data-based normalizations. Here, cell lines with similar morphologies show that individual PE or TG species scale similarly within their class, but not between classes. Two approaches are therefore compared for PQN, where the dilution factors were based on 1) all detected lipids or 2) lipids belonging to classes PE, PS, PI, PC and SM, the main components of plasma membranes in cells(van Meer & de Kroon, 2011).

When normalizing using all lipid abundances, PQN normalization is biased towards the large class of highly abundant TG species. For example, before normalization, class averages of TG and PE correlate moderately, due to similar overall amount of sample being measured (r = 0.65, **Figure 5A**). After normalization using all lipids, class averages of TG and PE per sample no longer show the positive correlation observed in the raw data but instead show a strong, artificial negative correlation (r = -0.76, **Figure 5B**). After normalization using only structural lipids, the positive trend between TG and PE is partially recovered (**Figure 5C**, r = 0.31).

**Figure 5.**
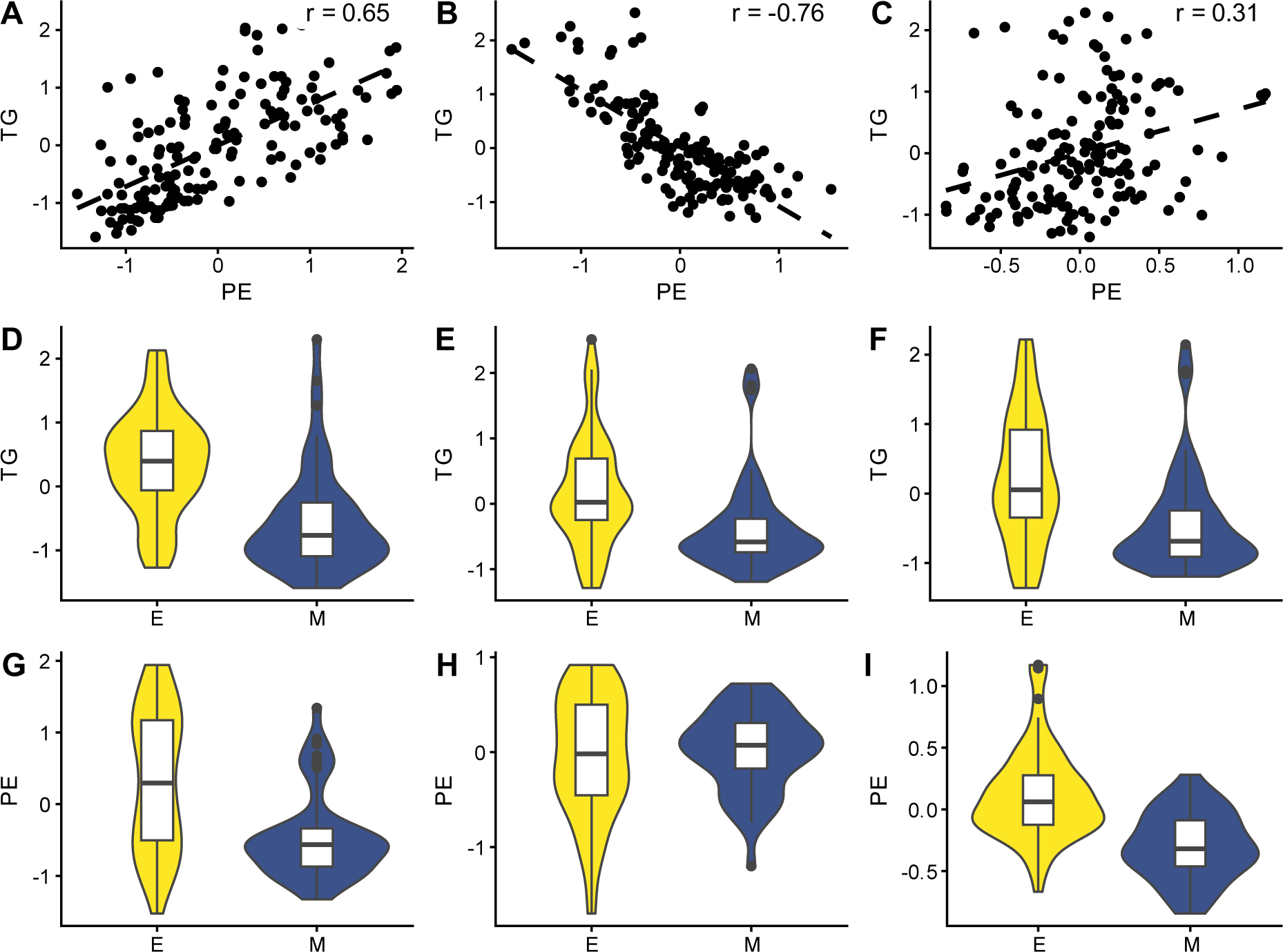
Differences in PE class average between morphologies are partially created by normalization. (A-C) (A) Lipid class averages for TG and PE before normalization of samples show a positive correlation (r=0.65). (B) After PQN normalization using all lipids, a negative correlation is observed (r=-0.76). (C) After PQN normalization with only structural lipids, the previously observed correlation is rescued to an extent (r=0.31). (D-I) (D) TG class averages are overall lower in mesenchymal cell lines before normalization, and after normalization using (E) all lipids or (F) only structural lipids. (E) In contrast, PE is lower in mesenchymal cells before normalization, but (H) similar after normalization using all lipids. (I) PQN using structural lipids does not artificially increase PE abundance. Class averages are the mean per class of log_2_ transformed autoscaled lipid abundances. E=epithelial, M=mesenchymal. Correlations are Pearson correlations.

The consequence of this bias towards TG was that PE abundances between mesenchymal and epithelial cell lines were flipped during PQN normalization. Before normalization, samples containing epithelial cell lines had overall higher PE and TG than mesenchymal cell lines (**Figure 5D, G**). However, we want to emphasize that this is likely due to the overall median signal per sample, as mentioned before (**Figure 1A** and **C**). When all data is used for normalization, mesenchymal cell lines had lower TG and similar PE compared to epithelial cell lines (**Figure 5E, H**). However, for the dataset normalized by structural lipids, both TG and PE were lower in mesenchymal cell lines (**Figure 5F, I**). With 347 detected triglycerides and only 65 phosphatidylethanolamines, this artificially higher PE abundance is caused by PQN and its underlying assumption that overall sample distributions must be the same. In summary, other lipid class averages are affected when cell lines have very different amounts of TGs and PQN normalization is used on all lipids.

### Log2 transformation and cube root transformation give comparable results

Transformation and scaling are also important parts of lipidomic data processing workflows. Lipidomic data is often log_2_ or cube root transformed before further analysis(van den Berg et al., 2006). Both log_2_ and cube root transformation increased ICC by a negligible amount compared to not transforming data ( **Supplementary Figure 2**). The resulting sample distributions showed right-skewed distributions for cube root transformed data and more symmetric distributions for log_2_ transformed data (**Supplementary Figure 3**).

To study if different transformations resulted in different outcomes, we compared how lipids contributed to the loadings of a PCA and performed a differential lipid analysis on a subset of the data. After PQN normalization with dilution factors based on structural lipids and either log_2_ or cube root transformation, followed by autoscaling, lipid classes showed similar contributions to the first three principal components (**Figure 6A-C**). Nine out of 10 lipids that contributed most to PC1 overlapped for log_2_ and cube root transformed data.

**Figure 6.**
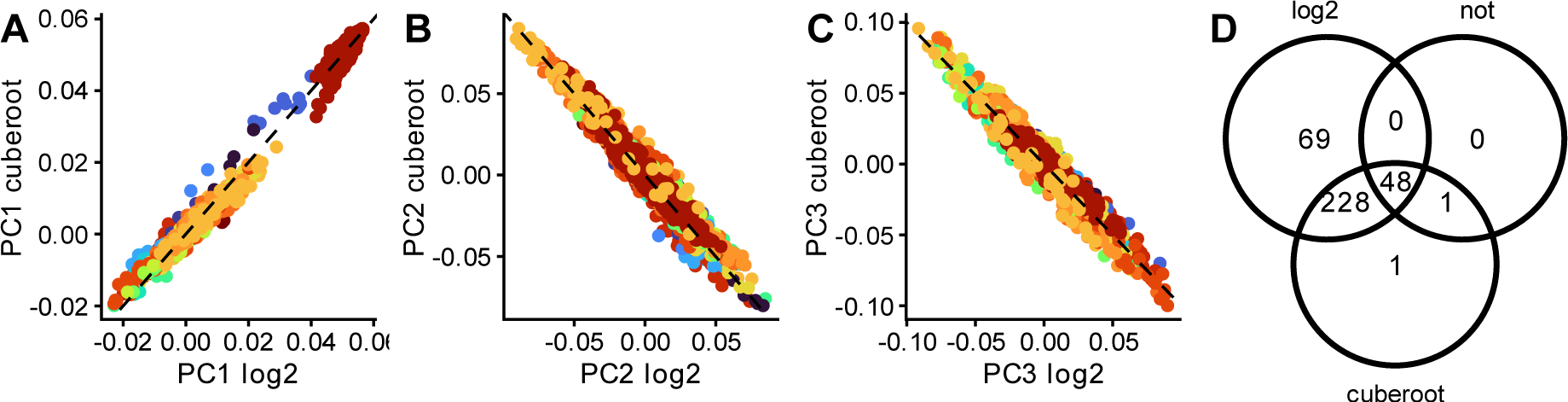
Correlation of principal components after data transformation shows similar results. (A-C) The (A) first, (B) second, and (C) third principal components after log_2_ or cube root transformation for lipids measured in cell lines show similar contributions of individual lipids. Black represents x=y. Each dot is the loading value of a lipid. Lipid classes are represented in different colors. Lipid abundances were scaled using autoscaling before performing PCA. (D) Overlap of differential abundant lipids between epithelial and mesenchymal morphologies without transformation or after log_2_ or cube root transformation.

Differential analysis is an alternative, univariate approach commonly used to study lipids with different abundances between groups. We calculated differential lipids between epithelial and mesenchymal cell lines using limma. Before transformation 49 lipids had different abundances (FDR < 0.1) while it was 278 or 345 after cube root or log_2_ transformation, respectively. The 69 additional hits after log_2_ were unique, and all other hits overlapped except 2 (**Figure 6D**). In conclusion, transforming data using log_2_ or cube root gave similar results.

## Discussion

In this study, we investigated different normalization techniques in lipidomic data based on either statistical metrics derived from the data itself or external biological parameters, such as cell count or protein concentration. While the latter would seem to be the ideal normalization strategy, it was shown here as well as in previous literature (Chao et al., 2017; Silva et al., 2013) to have the potential to introduce noise and thus obscure the signal caused by biological changes. However, a limitation for the data-based normalizations was also identified: a bias towards classes with numerous and highly abundant lipid species.

Normalization by protein concentration was insufficient to increase the correlation between replicates. Previous literature showed that normalization by protein concentration can give a relatively high error rate which was attributed to the preprocessing, such as washing and centrifuging steps, in a protein concentration assay (Chao et al., 2017). It is thus highly plausible that these steps also caused the low relation between lipid abundance per sample and protein concentration observed in this study. While less processing is needed for measuring cell counts, we observed that the cell counter sometimes counts clumps of epithelial cells as single cells (unpublished observation). This can cause some inaccuracy when cells clump together, which is more common for epithelial than for mesenchymal cell lines. In addition, cells in cell line panels often have different sizes (Dolfi et al., 2013; Ortmayr et al., 2019), which gives overall different lipid abundances per cell. This makes cell count less suitable for large-scale comparisons between cell line panels with different morphologies, as assayed here.

The underlying assumption for data-based normalization methods is that the overall distribution of lipids is the same across samples. However, we showed that lipid distributions and class averages varied depending on the chosen set of lipids used to normalize. This suggested an issue with lipids in the classes, particularly the class of triglycerides comprising numerous, highly abundant, and strongly correlating lipids. This bias cannot be detected using the ICC, because the ICC only indicates if replicates appear more similar after normalization. While more data-based approaches exist to normalize omics data, with advantages and limitations as compared previously by others (Chawade et al., 2014; Ejigu et al., 2013; Li et al., 2017; van den Berg et al., 2006), to our knowledge all of them are sensitive to large overall differences in sample content, and none provide a solution when a large class of lipids changes.

Triglycerides are components of lipid droplets, which concentrations can be much more dynamic than structural components of cell lines and serve as energy storage (Olzmann & Carvalho, 2019; Zadoorian et al., 2023). Sysi-Aho *et al*. previously observed a bias like the results observed in cells in the study presented here. They observed very different overall TG levels in obese and lean mice and mentioned that normalization could artificially alter phospholipid levels when total signal is used for normalization (Sysi-Aho et al., 2007), and proposed a new normalization algorithm that used a mix of internal standards for data normalization. In our opinion, this approach still assumes that the amount of starting material is similar. A previous study chose to perform normalizations within classes, thus preventing the problem of highly abundant lipid classes but limiting themselves to comparisons only within that class (Kindt et al., 2018). This is why we suggest using a probabilistic quotient normalization approach but based on only structural lipids as an appropriate alternative, as the raw values of these lipid classes (PE, PS, PI, PC and SM) correlated well with each other (range of r = 0.75-0.95, **Figure 4**). In addition, a normalization by structural lipids can overcome the limitation that cells in cell line panels often have different sizes.

One of the risks in choosing and testing multiple normalization strategies before data analysis, as performed in this study, is method mining as warned against by van den Berg *et al*.(van den Berg et al., 2006). The authors rightly state that one should not choose the method that best fits the data or expectations. However, in this study, the low correlation between protein concentration or cell count measures, as well as the large differences in median signal per sample were significant hurdles in analyzing the data appropriately. We suggest that using an independent metric, such as the ICC, is an appropriate middle between method mining and checking the quality of normalization.

Thorough quality checks are a must when choosing an appropriate normalization method, especially when dealing with lipidomic data. If highly similar replicates and cell count measurements are available, then using the ICC is recommended to check if normalization improves the agreement between replicates. After normalization, it is recommended to check the sample distributions, as aberrant class effects can influence any chosen normalization strategy and introduce noise into the results.

In conclusion, we show that biology-based normalizations cannot correct for all sample-specific variation in lipidomic data, especially when cell lines with different morphologies are investigated. Data-based normalizations, such as median or PQN normalization, improve similarity between replicates, but create artifacts if a large class with highly abundant lipids deviates. This is corrected by using dilution factors based on structural lipids. By providing this comparison of lipidomic normalization strategies, we hope to assist other scientists in choosing the appropriate normalization technique for investigating biological phenomena in large-scale cell line panels.

## Acknowledgments

We would like to thank Adel al Saaidy and Sabine Bos at the BioMedical Metabolomics Facility Leiden for their technical assistance with LC-MS/MS. This work was supported by the Netherlands Organization for Scientific Research (NWO) Enabling Technology Hotels program (project number: 435004026) and the China Scholarship Council (201608140084 and 201506220181). This publication is part of the “Building the infrastructure for Exposome research: Exposome-Scan” project (project number 175.2019.032) and EXPOSOME-NL funded through the Gravitation program of the Dutch Ministry of Education, Culture, and Science and the NWO (grant number 024.004.017).

---

**Supplementary Figure 1.**
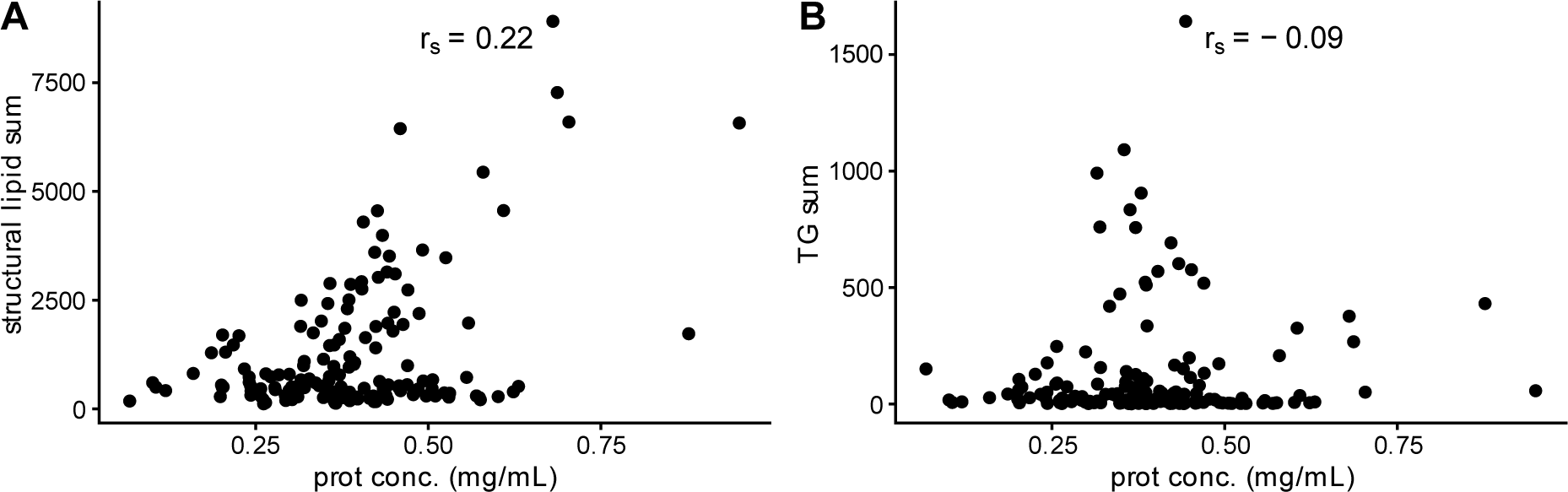
Structural lipids sometimes scale with protein concentration, but triglycerides do not. (A-B) sum of (A) all structural lipids or (B) triglycerides as a function of protein concentration. Lipids abundances are not transformed or scaled. r*_s_*: Spearman correlation coefficient.

**Supplementary Figure 2.**
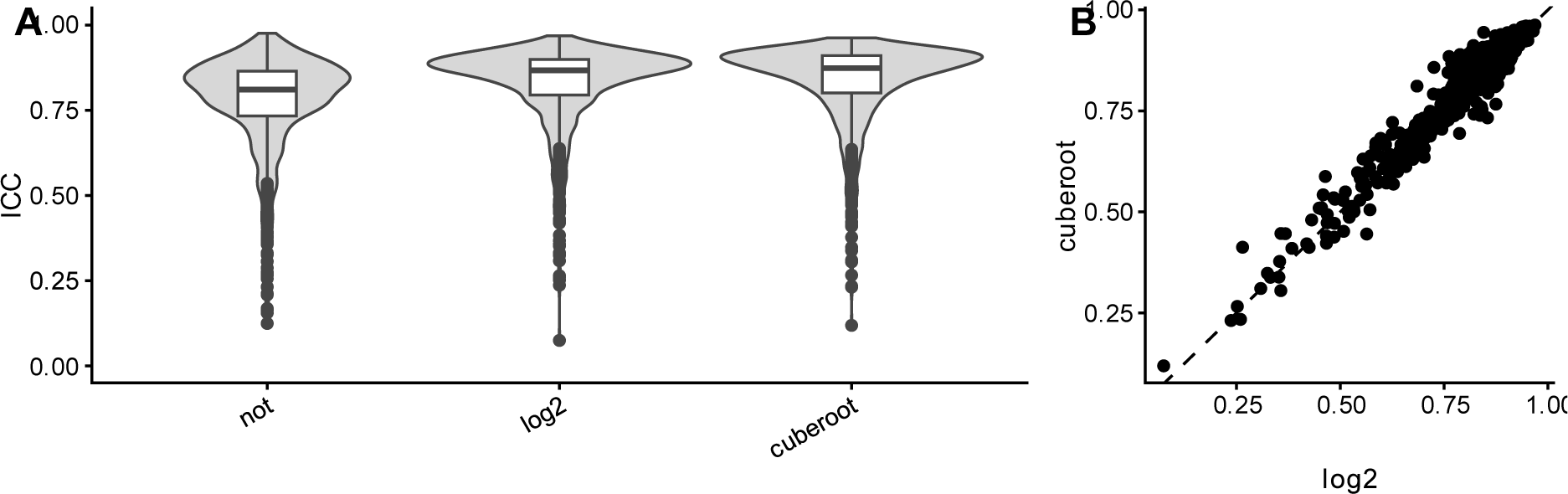
Distribution of ICC does not increase after log_2_ or cube root transformation. (A) ICC distributions before and after transformation. (B) ICC values for each lipid are similar after cube root or log_2_ transformation. Each dot is the ICC value of a lipid.

**Supplementary Figure 3.**
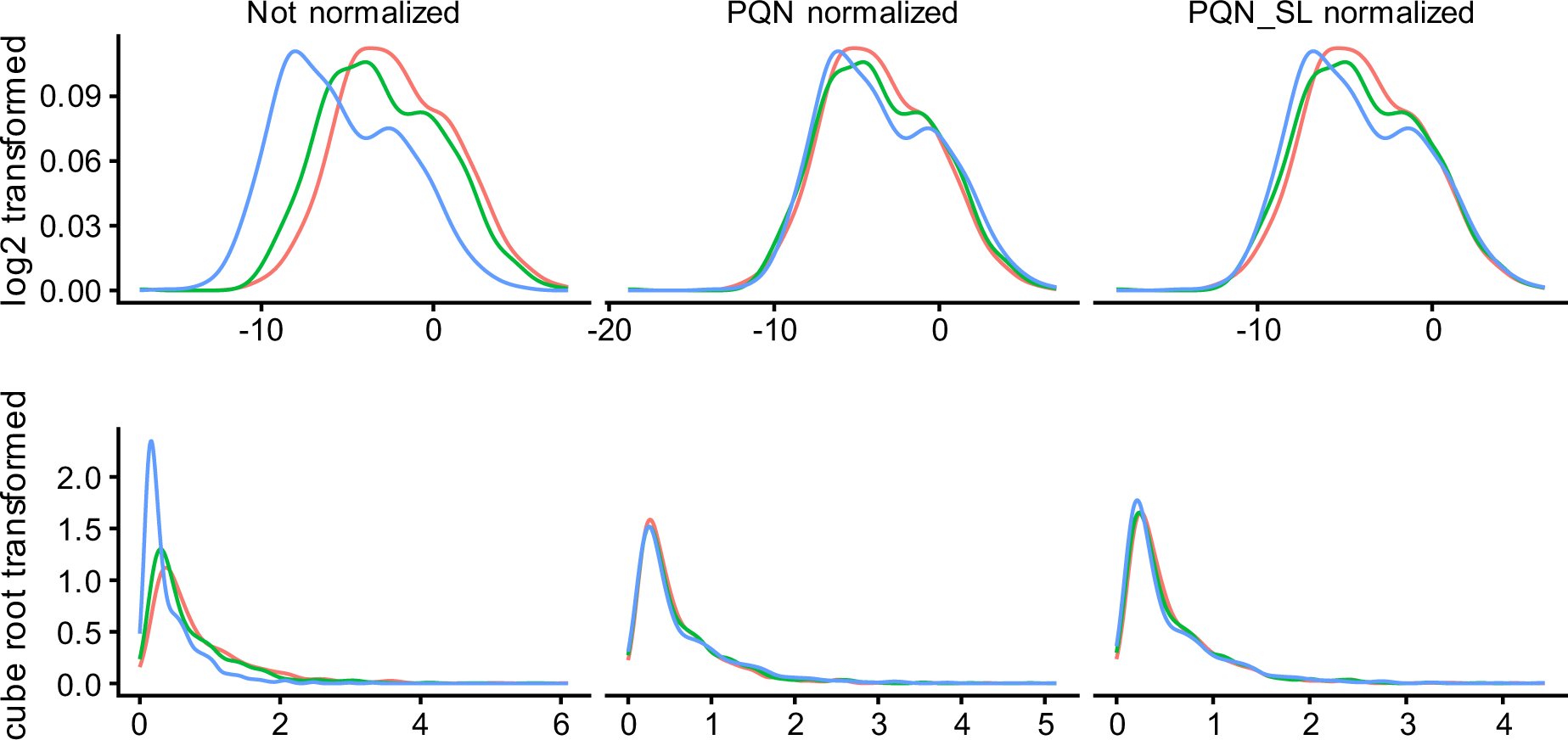
Shapes of distributions after normalization and transformation methods. Distributions of lipid abundance before normalization or after PQN normalization based on all lipids or only structural lipids. Colors represent the three replicates.

**Supplementary Table 1.** cell line morphology used in this project and the reference on which this is based.

| Cell_line | Cellosaurus_RRID | Morphology | Morphology_based_on |
| --- | --- | --- | --- |
| BT20 | CVCL_0178 | Epithelial-like | Rogkoti *et al.* (in preparation) |
| BT474 | CVCL_0179 | Epithelial-like | Rogkoti *et al.* (in preparation) |
| BT483 | CVCL_2319 | Epithelial-like | ATCC image, https://www.atcc.org/products/htb-121 |
| BT549 | CVCL_1092 | Mesenchymal-like | Rogkoti *et al.* (in preparation) |
| CAMA-1 | CVCL_1115 | Unknown |  |
| DU4475 | CVCL_1183 | Suspension | ATCC image, https://www.atcc.org/products/htb-123 |
| EVSA-T | CVCL_1207 | Epithelial-like | https://www.dsmz.de/collection/catalogue/details/culture/ACC-433 |
| HCC1143 | CVCL_1245 | Mesenchymal-like | Rogkoti *et al.* (in preparation) |
| HCC1187 | CVCL_1247 | Mesenchymal-like | ATCC image, https://www.atcc.org/products/crl-2322 |
| HCC1395 | CVCL_1249 | Mesenchymal-like | Rogkoti *et al.* (in preparation) |
| HCC1419 | CVCL_1251 | Epithelial-like | Rogkoti *et al.* (in preparation) |
| HCC1500 | CVCL_1254 | Epithelial-like | ATCC image, https://www.atcc.org/products/crl-2329 |
| HCC1569 | CVCL_1255 | Epithelial-like | Rogkoti *et al.* (in preparation) |
| HCC1599 | CVCL_1256 | Suspension | Sanger Institute, https://cellmodelpassports.sanger.ac.uk/downloads |
| HCC1806 | CVCL_1258 | Mesenchymal-like | Rogkoti *et al.* (in preparation) |
| HCC1937 | CVCL_0290 | Mesenchymal-like | Rogkoti *et al.* (in preparation) |
| HCC1954 | CVCL_1259 | Epithelial-like | Rogkoti *et al.* (in preparation) |
| HCC202 | CVCL_2062 | Epithelial-like | Rogkoti *et al.* (in preparation) |
| HCC2218 | CVCL_1263 | Suspension | Rogkoti *et al.* (in preparation) |
| HCC38 | CVCL_1267 | Mesenchymal-like | Rogkoti *et al.* (in preparation) |
| HCC70 | CVCL_1270 | Mesenchymal-like | Rogkoti *et al.* (in preparation) |
| Hs578T | CVCL_0332 | Mesenchymal-like | Rogkoti *et al.* (in preparation) |
| MCF-7 | CVCL_0031 | Epithelial-like | Rogkoti *et al.* (in preparation) |
| MDA-MB-134VI | CVCL_0617 | Round | Rogkoti *et al.* (in preparation) |
| MDA-MB-157 | CVCL_0618 | Unknown |  |
| MDA-MB-175VII | CVCL_1400 | Epithelial-like | Rogkoti *et al.* (in preparation) |
| MDA-MB-231 | CVCL_0062 | Mesenchymal-like | Rogkoti *et al.* (in preparation) |
| MDA-MB-330 | CVCL_0619 | Epithelial-like | ATCC image, https://www.atcc.org/products/htb-127 |
| MDA-MB-361 | CVCL_0620 | Epithelial-like | Rogkoti *et al.* (in preparation) |
| MDA-MB-415 | CVCL_0621 | Epithelial-like | Rogkoti *et al.* (in preparation) |
| MDA-MB-436 | CVCL_0623 | Mesenchymal-like | Rogkoti *et al.* (in preparation) |
| MDA-MB-453 | CVCL_0418 | Round | Rogkoti *et al.* (in preparation) |
| MDA-MB-468 | CVCL_0419 | Round | Rogkoti *et al.* (in preparation) |
| MPE600 | CVCL_9875 | Unknown |  |
| OCUB-F | CVCL_3352 | Suspension | Rogkoti *et al.* (in preparation) |
| OCUB-M | CVCL_1621 | Round | Rogkoti *et al.* (in preparation) |
| SK-BR-3 | CVCL_0033 | Epithelial-like | ATCC image, https://www.atcc.org/products/htb-30 |
| SK-BR-5 | CVCL_5217 | Unknown |  |
| SK-BR-7 | CVCL_5218 | Mesenchymal-like | Rogkoti *et al.* (in preparation) |
| SUM102PT | CVCL_3421 | Mesenchymal-like | Rogkoti *et al.* (in preparation) |
| SUM1315MO2 | CVCL_5589 | Mesenchymal-like | Rogkoti *et al.* (in preparation) |
| SUM149PT | CVCL_3422 | Mesenchymal-like | Rogkoti *et al.* (in preparation) |
| SUM159PT | CVCL_5423 | Mesenchymal-like | Rogkoti *et al.* (in preparation) |
| SUM185PE | CVCL_5591 | Epithelial-like | Rogkoti *et al.* (in preparation) |
| SUM190PT | CVCL_3423 | Epithelial-like | Rogkoti *et al.* (in preparation) |
| SUM225CWN | CVCL_5593 | Unknown |  |
| SUM229PE | CVCL_5594 | Mesenchymal-like | Rogkoti *et al.* (in preparation) |
| SUM44PE | CVCL_3424 | Round | Rogkoti *et al.* (in preparation) |
| SUM52PE | CVCL_3425 | Epithelial-like | Rogkoti *et al.* (in preparation) |
| T47D | CVCL_0553 | Epithelial-like | Rogkoti *et al.* (in preparation) |
| UACC812 | CVCL_1781 | Epithelial-like | Rogkoti *et al.* (in preparation) |
| UACC893 | CVCL_1782 | Epithelial-like | Rogkoti *et al.* (in preparation) |
| ZR75-1 | CVCL_0588 | Epithelial-like | Rogkoti *et al.* (in preparation) |
| ZR75-30 | CVCL_1661 | Epithelial-like | ATCC image, https://www.atcc.org/products/crl-1504 |

